# *Desulfovibrio diazotrophica* sp. nov., a sulphate reducing bacterium from the human gut capable of nitrogen fixation

**DOI:** 10.1101/2020.07.01.183566

**Authors:** Lizbeth Sayavedra, Tianqi Li, Marcelo Bueno Batista, Brandon K.B. Seah, Catherine Booth, Qixiao Zhai, Wei Chen, Arjan Narbad

## Abstract

Sulphate-reducing bacteria (SRB) are widespread in human guts, yet their expansion has been linked to colonic diseases. We report the isolation, genome sequencing, and physiological characterisation of a novel SRB species belonging to the class *Deltaproteobacteria* (QI0027^T^). Phylogenomic analysis revealed that the QI0027^T^ strain belongs to the genus *Desulfovibrio* with its closest relative being *Desulfovibrio legallii*. Metagenomic sequencing of stool samples from 45 individuals, as well as comparison with 1690 *Desulfovibrionaceae* metagenome-assembled genomes, revealed the presence of QI0027^T^ in at least 22 further individuals. QI0027^T^ encoded nitrogen fixation genes and based on the acetylene reduction assay, actively fixed nitrogen. Transcriptomics revealed that QI0027^T^ overexpressed 45 genes in nitrogen limiting conditions as compared to cultures supplemented with ammonia, including nitrogenases, an urea uptake system and the urease enzyme complex. To the best of our knowledge, this is the first *Desulfovibrio* human isolate for which nitrogen fixation has been demonstrated. This isolate was named *Desulfovibrio diazotrophica* sp. nov., referring to its ability to fix nitrogen (‘diazotroph’).

**Importance:** Animals are often nitrogen limited and have evolved diverse strategies to capture biologically active nitrogen. These strategies range from amino acid transporters to stable associations with beneficial microbes that can provide fixed nitrogen. Although frequently thought as a nutrient-rich environment, nitrogen fixation can occur in the human gut of some populations, but so far it has been attributed mainly to *Clostridia* and *Klebsiella* based on sequencing. We have cultivated a novel *Desulfovibrio* from human gut origin which encoded, expressed and actively used nitrogen fixation genes, suggesting that some sulphate reducing bacteria could also play a role in the availability of nitrogen in the gut.

## Introduction

Sulphate reducing bacteria (SRB) are present in the mouth and the gut of ~50% of the human population (1, 2). SRB thrive in the gut, releasing hydrogen sulphide (H_2_S) as a by-product of sulphate reduction. H_2_S is a potent genotoxin and has been linked to chronic colonic disorders and inflammation of the large intestine (3). Likewise, the presence of some *Desulfovibrio* species has been implicated in chronic periodontitis, cell death and inflammatory bowel diseases such as ulcerative colitis and Crohn’s disease (4). The detrimental role of SRB is, however, not firmly established. H_2_S can also act as a signalling molecule or energy source for mitochondria (5, 6). Moreover, by using hydrogen, SRB help in the efficient energy acquisition and complete oxidation of substrates produced by fermentative bacteria (7). SRB could, therefore, have a dual role in the gut microbiome.

Relatively few strains from healthy host individuals have been characterized. Cultivated SRB isolated from humans include *Desulfovibrio piger*, *D. fairfieldensis*, *D. desulfuricans* and *D. legallii* (4, 8), with *D. piger* described as the most abundant in samples obtained from western countries (9). The physiology and genomic potential of strains isolated from non-western countries remain largely unexplored.

Biological nitrogen fixation is the process by which gaseous dinitrogen (N_2_) is reduced to biologically available ammonia (NH_3_) by diazotrophic microbes (10). Diazotrophy was reported in some *Desulfovibrio* species from free-living communities and the termite gut (11, 12). All animals require access to a source of fixed nitrogen (13). Most of the readily available nitrogen in humans comes from food, but nitrogen fixation can occur in the microbiome of some human populations (14). In the human microbiome, the potential to fix nitrogen has been so far linked to *Klebsiella* and *Clostridia* (14). The human host can regulate the nitrogen available to the microbiome through the diet and intestinal secretions (15), so bacteria that can fix nitrogen could potentially have a competitive advantage in the gut environment.

We report here the isolation and characterization of a novel species of the genus *Desulfovibrio*, strain QI0027^T^, isolated from a stool sample provided by a Chinese male donor. Surprisingly this strain encoded the genes for nitrogen fixation. The goals of this study were (i) to characterize strain QI0027^T^, (ii) to identify if diazotrophy is commonly distributed among *Desulfovibrionaceae* associated to humans and (iii) to examine if the genes required for nitrogen fixation are expressed and functional. We used metagenomic sequencing of 45 human donors, as well as analysed comprehensive collections of genomes recovered from metagenomes to examine its distribution. We used physiological tests to examine the nitrogenase activity. Finally, we used transcriptomics to determine the genes that are expressed by QI0027^T^ in response to the absence of a source of fixed nitrogen.

## Results

### Genome sequencing and phylogenomics

As part of a study to isolate and characterize SRB from Chinese donors, we isolated strain QI0027^T^. Sequencing of QI0027^T^ resulted in a 2.85 Mb genome, which was 99.1% complete based on conserved *Deltaproteobacteria* marker genes and had a 62.1% GC content (Table 1). Comparison of the 16S rRNA from QI0027^T^ against the SILVA SSU r138 database revealed that the isolate belongs to the *Deltaproteobacteria* class and that its closest relative is *Desulfovibrio legallii*. Even though the 16S rRNA gene of QI0027^T^ shared a 99% identity with the partial 16S rRNA sequence of *D. legallii*, the average nucleotide identity (ANI) of their whole genomes was only 86.49%, which was well below the recommended threshold for species discrimination of 94-96% ANI (16). Thus, QI0027^T^ and *D. legallii* are different species.

**Table 1.** Differentiation between *D. legallii* and QI0027^T^. S = sensitive; R = Resistant; +/− = Light resistance

|  |  | QI0027<sup>T</sup> | *D. legallii* |
| --- | --- | --- | --- |
| Genome | GC (%) content | 62.11 | 64.81 |
|  | Genome size (Mbp) | 2.85 | 2.7 |
|  | Genome completeness* (%) | 99.11 | 100 |
|  | Genome contamination (%) | 0.61 | 0.59 |
|  | Strain heterogeneity (%) | 0 | 0 |
| Physiology | Antibiotic resistance |  |  |
|  | Ampicillin (25 µg) | S | S |
|  | Chloramphenicol (10 µg) | S | S |
|  | Nalidixic acid (30 µg) | S | S |
|  | Kanamycin (30 µg) | R (+/-) | R (+/-) |
|  | Erythromycin (30 µg) | S | S |
|  | Tetracycline (30 µg) | S | S |
|  | Streptomycin (25 µg) | R | R |

\*Genome completeness, contamination, and strain heterogeneity was determined with CheckM based on highly conserved genes among Deltaproteobacteria.

Consistent with our 16S rRNA comparison, the closest relative of QI0027^T^ was *D. legallii,* and these bacteria grouped within the *‘sensu-stricto’ Desulfovibrio* based on a well-supported phylogenomic tree reconstructed using 30 marker genes (17) (Figure 1). The family *Desulfovibrionaceae* has eight described genera: *Desulfovibrio, Bilophila, Lawsonia, Halodesulfovibrio, Pseudodesulfovibrio, Desulfocurvus, and Desulfobaculum*, although no representative genomes have been sequenced for the former two genera. Bacteria from the genera *Maihella*, *Bilophila* and *Lawsonia* grouped closer to the ‘*sensu-stricto’ Desulfovibrio* as compared to *Halodesulfovibrio, Pseudodesulfovibrio,* and other *Desulfovibrio* bacteria that despite their names, are different genera from the *Desulfovibrionaceae* family (Figure 1) (18).

**Figure 1.**
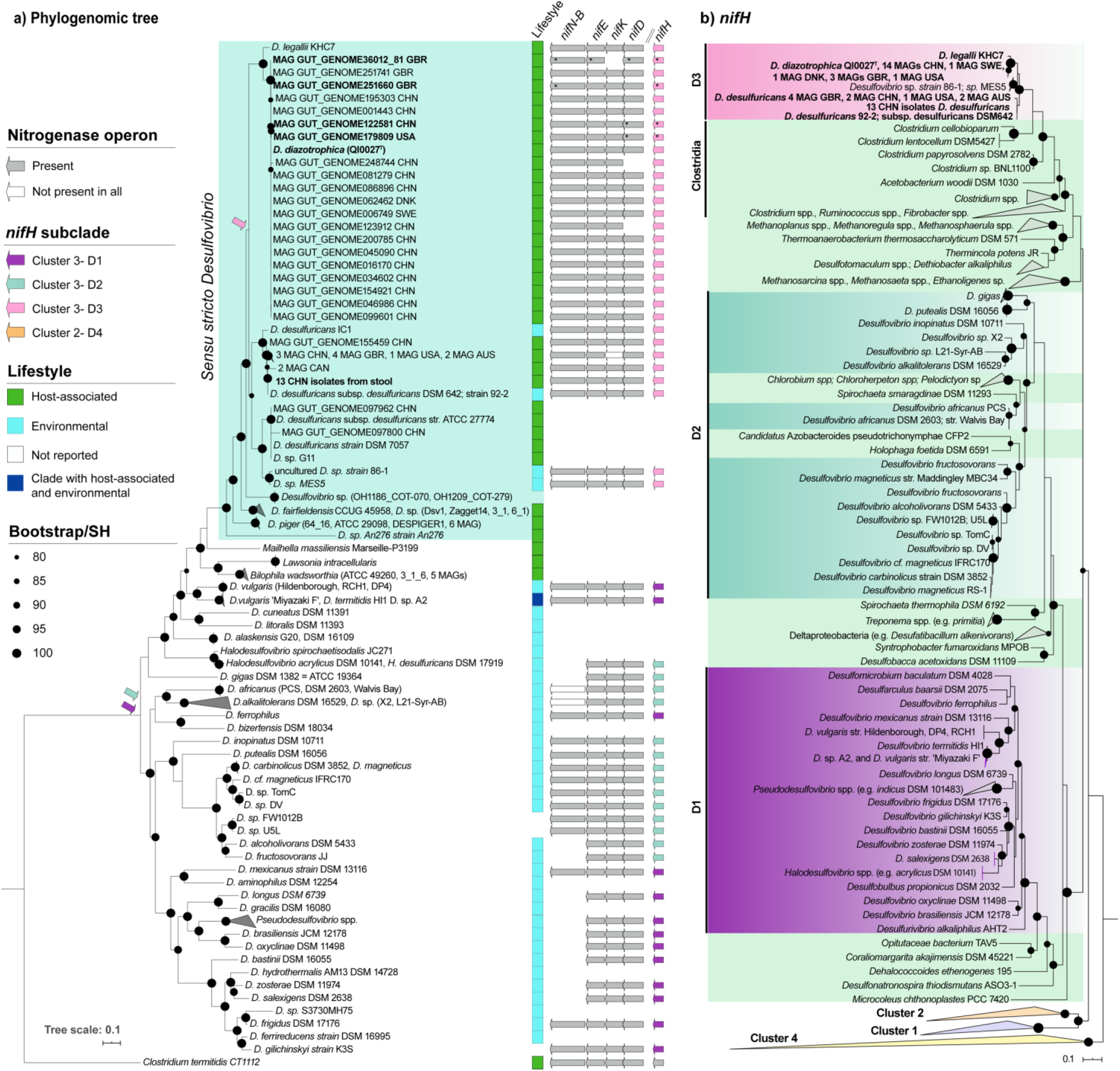
Phylogenomic tree reconstruction of *Desulfovibrionaceae* bacteria with gene presence/absence of the nitrogenase cluster (a) and *nifH* phylogeny (b). (a) Phylogenomic tree reconstruction revealed that strain (QI0027^T^) belongs to *Desulfovibrio* ‘*sensu-stricto*’ group. The maximum-likelihood tree was reconstructed using the amino acid sequences encoded by 30 single-copy marker genes conserved among the *Desulfovibrionaceae* bacteria extracted with Amphora2. The tree was rooted with *Clostridium termitidis* CT1112. Presence of genes encoding the Fe protein (also known as dinitrogenase reductase, *nifH*), the molybdenum-iron (Mo-Fe) protein (also known as dinitrogenase, *nifD* and *nifK*) and encoding proteins required for iron-molybdenum cofactor (FeMo-co) biosynthesis (*nifB* and *nifE*) is shown. Genes recovered using targeted assembly for QI0027^T^ MAGs are shown with *. Purple, blue and pink arrows on phylogenomic tree show when *nifH* was likely acquired. The country of origin from which gut metagenomes were recovered is shown with the ISO3 code. (b) Maximum likelihood of *nifH* gene product tree reconstruction. 581 amino acid sequences spanning 300 amino acid positions were used to reconstruct the phylogeny. Cluster 4 contains paralogous genes that are not involved in nitrogen fixation. Solid circles in phylogeny represent SH-like support between 0.8 -1.0, with the size proportional to the support value. Bootstrap support (a) or SH support (b) between 80-100% is represented as filled circles scaled to size

### Physiology

Similar to other *Desulfovibrionaceae* bacteria, QIOO27^T^ is a gram-negative bacterium with curved-rod to spirochete shape (Figure 2). Cells were 1.4 to 5 μm long. QI0027^T^ grew at a temperature between 15 to 46°C, with optimum growth temperature of 37°C. QI0027^T^ could grow with 0-0.51 M NaCl, with optimum growth at 0.068-0.171 M NaCl. This salinity concentration range is comparable with human physiological saline (0.9% w/v, 0.15 M) (19). Colonies on Postgate C plates were black due to the accumulation of iron sulphide precipitates (Supplementary Figure 1A). Based on disc diffusion assays with antibiotics, QI0027^T^ was resistant to kanamycin and streptomycin (Table 1).

**Figure 2.**
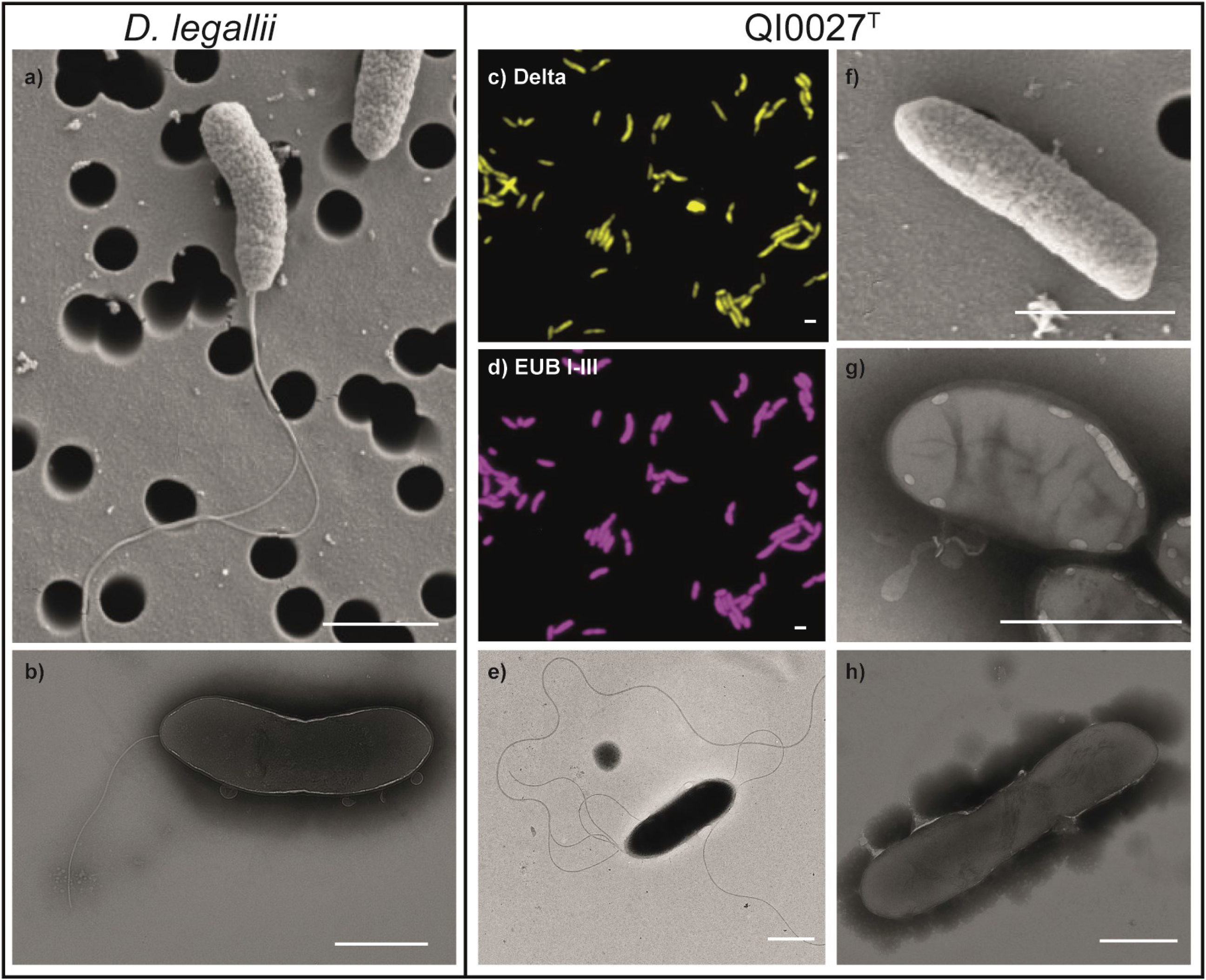
Comparative microscopic characteristics of *D. legallii* and QI0027^T^. Scanning electron microscopy (SEM) (a) and transmission electron microscopy (TEM) with negative staining (b) showed the presence of a single polar flagellum in *D. legallii*. Outer membrane vesicle-like structures could be observed in *D. legallii* (b). SEM (f) and TEM (g,h) show that most QI0027T cells did not have flagella, with a few exceptions (e). Some cells also showed inclusion bodies, which might be polyhydroxyalkanoates (g) (Hai *et al.*, 2004). Scale bar = 1 μm.

Unlike the clearly motile *D. legallii*, QI0027^T^ was mostly non-motile but showed occasionally multiple lophotrichous flagella based on a spatial assay on soft agar and scanning and transmission electron microscopy (SEM and TEM). When bacteria are non-motile, they will show a sharp edge where the inoculation stab is made. In contrast, if they are motile, they will show a cloudy growth around the stabbing site. *D. legallii* showed a diffuse area of growth, in agreement with the previously reported presence of a flagellum (8). In contrast, QI0027^T^ had a sharp edge area of growth. However, there were slight signs of iron sulphide outside the area of the stabbing site, which could be due to some cells being motile or the diffusion of the hydrogen sulphide gas produced by QI0027^T^ (Supplementary Figure 1B). To further investigate the presence of flagellum in QI0027^T^, we used SEM and negative staining combined with TEM. *D. legallii* showed a single polar flagellum with both EM techniques (Figure 2). SEM showed that QI0027^T^ cells did not have a flagellum. However, while TEM showed that most cells did not have a flagellum, we could observe occasionally some cells with multiple lophotrichous flagella. The flagella of QI0027^T^ were longer and thinner as those compared to *D. legallii*, which might explain why we could not observe them with SEM, as the sample processing is harsher on the cells as compared to the negative staining. QI0027^T^ encoded the genes for flagella synthesis, including the hook-basal body complex (*fliE*), the basal-body rod (*flgB, flgC*), the cytoplasmic components *fliI*, and the membrane-bound components (*fliO, fliP, fliQ, flhA, flhB, fliR*) of the flagellar protein export apparatus (reviewed by (20)). QI0027^T^ also synthesized the negative regulator of flagellin synthesis *flgM*. We thus hypothesize that QI0027^T^ regulates the expression of flagella synthesis, which might be a key factor during the colonization and sensing of areas rich in nutrients in the gastrointestinal tract (21).

Fatty acid analysis from QI0027^T^ and *D. legallii* showed that the main fatty acid was iso C17:1 (29% for QI0027^T^ and 24.2% for *D. legallii*), which is consistent with most *Desulfovibrio* species (22). Other predominant fatty acids had also a similar composition and abundance for both strains: iso fatty acid methyl ester (FAME) C_17:0_ (24.9% for QI0027^T^ and 26.12% for *D. legallii*), FAME C_15:0_ (19.3% for QI0027^T^ and 20.6% for *D. legallii*)), and iso C_17:0_ (17.2-17.5% for QI0027^T^ and 19.9-20.2% for *D. legallii*) (Table 2). However, 10 fatty acids were detected at low abundance (< 2%) only in QI0027^T^, which likely reflects small differences in the metabolism of QI0027^T^ compared to *D. legallii* (Table 2).

**Table 2.** Cellular fatty acid composition (%) of QI0027^T^ and its closest relative *D. legallii* (DSM 19129) grown in Postgate C media. FAME, fatty acid methyl ester; ND, not detected

| Calculation method | Cellular fatty acids | QI0027<sup>T</sup> | *D. legallii* |
| --- | --- | --- | --- |
| TSBA40 | iso C17:1ω9c | 29.01 | 24.22 |
| ANAER6 | iso 17:0 FAME | 24.94 | 26.12 |
| ANAER6 | iso 15:0 FAME | 19.30 | 20.66 |
| TSBA6 | iso 17:0 | 17.53 | 20.20 |
| ANAER6 | 16:0 FAME | 15.01 | 16.53 |
| TSBA6 | iso 15:0 | 13.56 | 15.98 |
| ANAER6 | 18:0 FAME | 11.25 | 9.92 |
| TSBA6 | 16:0 | 10.55 | 12.78 |
| TSBA6 | 18:0 | 7.90 | 7.67 |
| ANAER6 | anteiso 15:0 FAME | 5.28 | 8.06 |
| ANAER6 | 16:1 cis 9 FAME | 3.87 | 3.57 |
| TSBA6 | 15:0 anteiso | 3.71 | 6.23 |
| ANAER6 | 16:0 dimethyl acetal | 3.32 | 3.02 |
| ANAER6 | 17:0 cyc FAME | 3.04 | 1.66 |
| ANAER6 | UN 17.103 17:0i dimethyl acetal | 2.97 | 2.86 |
| TSBA6 | Sum in feature 3 - 16:1 ω7c/16:1 ω6c and/or 16:1 ω6c/16:1 ω7c | 2.72 | 2.76 |
| TSBA40 | Sum in feature 3 - 16:1 ω7c/15 iso 2OH | 2.67 | 2.73 |
| ANAER6 | 16:0 aldehyde | 2.40 | 1.46 |
| TSBA6 | Sum in feature 4 - 17:1 iso I/anteiso B | 2.33 | 2.34 |
| ANAER6 | 16:0 iso FAME | 2.16 | 3.40 |
| TSBA6 | 17:0 cyclo | 2.13 | 1.28 |
| ANAER6 | 17:0 anteiso FAME | 1.94 | 2.73 |
| ANAER6 | 15:0 iso dimethyl acetal | 1.90 | ND |
| ANAER6 | 12:0 FAME | 1.71 | ND |
| TSBA40 | unknown 14.959 | 1.66 | 1.12 |
| TSBA6 | 16:0 iso | 1.52 | 2.63 |
| TSBA6 | 16:1 ω7c alcohol | 1.45 | ND |
| TSBA6 | anteiso 17:0 | 1.36 | 2.11 |
| TSBA6 | anteiso 17:1 ω9c | 1.35 | 1.52 |
| TSBA40 | anteiso 17:0 | 1.34 | 2.09 |
| TSBA6 | iso 14:0 3OH | 1.33 | ND |
| TSBA40 | anteiso 17:1 ω9c | 1.33 | 1.51 |
| TSBA40 | iso 14:0 3OH | 1.31 | ND |
| TSBA6 | 12:0 | 1.20 | ND |
| TSBA6 | iso 16:1 H | 1.19 | ND |
| TSBA40 | 12:0 | 1.18 | ND |
| ANAER6 | 10:0 FAME | 0.94 | ND |
| TSBA6 | 10:0 | 0.66 | ND |

We used a microplate system, which allows for the high-throughput monitoring of 96 carbon substrates under anaerobic conditions, to identify the carbon substrates used by QI0027^T^ and *D. legallii* (Biolog, Technopath, Ireland). Substrates that could be used by both, QI0027^T^ and *D. legallii* strains, included L-rhamnose, D-fructose, L-fucose, D-galactose, D-galacturonic acid, palatinose, D,L-lactic acid, L-lactic acid, D-lactic acid methyl ester, L-malic acid, pyruvic acid, methyl pyruvate, and L-glutamic acid, glucose and D-mannose (Table 3). Only QI0027^T^ used as carbon substrate B-gentiobiose, D-glucose-6-phosphate, and D-melibiose. Likewise, only *D. legallii* used arbutin, α-keto-butyric acid, α -ketovaleric acid, L-methionine, L-valine, and uridine-5’-monophosphate. However, replicates of each species did not show consistent use of these carbon substrates, suggesting that their utilization might not be a reliable method to distinguish between the two bacterial species.

**Table 3.** Carbon compounds used by *D. legallii* and QI0027^T^ as determined with the AN microplate system (Biolog). The use of each carbon substrate was assigned as being used or not based on the function do_disc from the opm package, using k-means, with no weak reactions for each of three replicates.

| Carbon source | Carbon compound used?<br>(N=3) |  |
| --- | --- | --- |
|  | *D. legallii* | QI0027<sup>T</sup> |
| Arbutin | 1/3 | 0/3 |
| B-cyclodextrin | 1/3 | 1/3 |
| Dextrin | 1/3 | 1/3 |
| D-fructose | 2/3 | 2/3 |
| L-fucose | 3/3 | 2/3 |
| D-galactose | 3/3 | 2/3 |
| D-galacturonic acid | 2/3 | 2/3 |
| B-Gentiobiose | 0/3 | 2/3 |
| D-glucose | 2/3 | 1/3 |
| D-Glucose-6-Phosphate | 0/3 | 1/3 |
| D-Mannose | 2/3 | 1/3 |
| D-Melibiose | 0/3 | 1/3 |
| 3-O-Methyl-D-Glucose | 1/3 | 1/3 |
|  | *D. legallii* | QI0027<sup>T</sup> |
| Palatinose | 2/3 | 2/3 |
| L-Rhamnose | 3/3 | 3/3 |
| Turanose | 1/3 | 1/3 |
| Acetic acid | 1/3 | 1/3 |
| Formic acid | 1/3 | 1/3 |
| Glyoxylic acid | 1/3 | 1/3 |
| A-Hydroxybutyric acid | 1/3 | 1/3 |
| A-keto-butyric acid | 1/3 | 0/3 |
| A-ketovaleric acid | 1/3 | 0/3 |
| D,L-Lactic acid | 1/3 | 2/3 |
| L-lactic acid | 1/3 | 2/3 |
| D-lactic acid methyl ester | 2/3 | 2/3 |
| L-malic acid | 3/3 | 2/3 |
| Propionic acid | 1/3 | 1/3 |
| pyruvic acid | 1/3 | 2/3 |
| methyl pyruvate | 3/3 | 2/3 |
| urocanic acid | 1/3 | 1/3 |
| Alaninamide | 1/3 | 1/3 |
| L-glutamic acid | 2/3 | 2/3 |
| L-methionine | 1/3 | 0/3 |
| L-serine | 1/3 | 1/3 |
| L-valine | 1/3 | 0/3 |
| L-valine plus L-Aspartic acid | 1/3 | 1/3 |
| Uridine-5'-monophosphate | 1/3 | 0/3 |

### General metabolic reconstruction

Based on our genome reconstruction, QI0027^T^ can reduce sulphate to hydrogen sulphide, using either hydrogen, carbon monoxide or alcohols as electron donors (Supplementary Figure 2). For the oxidation of alcohols, an alcohol dehydrogenase could be used for the conversion of ethanol to acetate, similar to *D. gigas* (Kremer *et al.*, 1988). Lactate could be oxidized to pyruvate by a membrane-bound lactate dehydrogenase. Ammonia could be further used for the biosynthesis of glutamine, glutamate, asparagine, NAD and deazapurine (Supplementary Figure 2). QI0027^T^ encoded the genes for the amino acid biosynthesis of methionine, asparagine, alanine, arginine, threonine, isoleucine, aspartate, serine, glycine, leucine, and valine. Additionally, QI0027^T^ could synthesize the following vitamins and cofactors: biotin (B7), folate (B9), phosphopantothenate (B5), thiamine (B1), pyridoxal (B6), adenosylcobalamin (B12) and flavin.

### Distribution of QI0027-related species in human microbiome metagenomes

We detected the presence of the same species as QI0027^T^ on at least two further individuals based on sequencing of 45 stool metagenomes from Chinese donors. We used three methods to detect QI0027^T^ in these Chinese metagenomes. First, we used mOTUS2, a taxonomic classifier that uses clade-specific marker genes (23). The whole-genome sequencing (WGS) reads from the pure culture QI0027^T^ were used as a control to identify the assigned species by the taxonomic classifier. mOTUS2 classified the WGS reads only at genus level as *Desulfovibrio*. By including in the mOTUS2 reference database our QI0027^T^ genome assembly, mOTUS2 was able to detect QI0027^T^ in one individual. Second, we used MetaPhlAn2 (24), a taxonomic classifier that uses a different database of clade-specific marker genes. MetaPhlAn2 is widely used in studies that combine metagenomic sequencing efforts from diverse human microbiome studies (e.g. 25, 26). MetaPhlAn2 classified the reads of QI0027^T^ as ‘*Desulfovibrio desulfuricans’*, even though the closest relative is the type strain *D. legallii*. Reads classified as ‘*D. desulfuricans*’ by MetaPhlAn2 will therefore include other species such as QI0027^T^. The lack of specificity or mis classification of QI0027^T^ likely reflects the lack of representative genomes in the MetaPhlAn2 database. Third, we used read mapping against our QI0027^T^ genome assembly using a high similarity threshold to overcome the identification limitation by these taxonomic profilers. Using this method, we identified the presence of QI0027^T^ in two individuals (Supplementary Dataset 1).

Metagenomic reads from the stool donor that was used for isolating QI0027^T^ only covered 0.1% of the reference genome and had an average fold coverage of 0.0082. Overall, read mapping using QI0027^T^ genome as reference had a higher sensitivity to detect this species in the gut microbiome as compared to mOTUs2, and a lower rate of false positives as compared to MetaPhlAn2.

Based on an ANI similarity of 99.49%, we successfully recovered a high-quality draft genome of QI0027^T^ species from metagenomic reads, based on a targeted assembly approach of *Desulfovibrionaceae* using the 45 stool metagenomes from Chinese donors. This targeted assembly approach was used to evaluate if the genome of this species could be recovered in high-complexity stool metagenomes. Surprisingly, we were not able to recover *D. desulfuricans*, even though this was the species that could be isolated more often with our isolation efforts in Chinese stool samples (13 out of 14 *Desulfovibrionaceae* isolates). Our results suggest that higher sequencing depth would have been necessary to confidently detect the full species diversity of SRB using metagenomic approaches and that culturing approaches might reflect a different picture of the SRB diversity.

We further identified 20 genomes from the same species as QI0027^T^ (ANI > 98%) using large unified collections of metagenome-assembled genomes (MAGs) obtained from extensive diversity of lifestyles, ages and countries (1, 25). The genome collection (25) included 1690 *Desulfovibrionaceae* genomes classified as *D. piger, D. desulfuricans*, *D. fairfieldensis, Desulfovibrio* sp*., Bilophila* sp., *B. wadsworthia,* and *Maihella* sp. Genomes from the same species as QI0027^T^ were assembled from stool metagenomes obtained from 14 donors residing in China, three in UK, one in Sweden, one in Denmark, and one in USA (Metagenome studies from 27, 28–32). Since genomes of organisms that are more abundant are easier to recover from metagenomes (33), the distribution of QI0027^T^ among humans from diverse countries suggests that this species is more prevalent in Chinese individuals.

### QI0027^T^ encoded, expressed and actively used nitrogen fixation genes

Genome analysis of the strain QI0027^T^ revealed the presence of the minimal gene set for nitrogen fixation (34). These genes included the key gene *nifH*, encoding the dinitrogenase reductase; *nifD* and *nifK,* encoding the MoFe dinitrogenase; as well as *nifE*, *nifN* and *nifB*, encoding the FeMo cofactor biosynthesis machinery. The *nifN* and *nifB* genes were fused into a single gene (*nifN-B*). Two genes encoding PII-like nitrogen regulatory proteins (*glnB1* and *glnB2*) were encoded between *nifH* and *nifD* as observed in *Clostridium* and other diazotrophic *Desulfovibrio* (Supplementary Figure 3) (10, 35). To physiologically validate the nitrogen fixation ability of strain QI0027^T^, we checked the expression pattern of the genes encoding the nitrogen fixation machinery under conditions of nitrogen limitation compared to nitrogen excess. As expected, we observed that all genes in the *nif* operon were significantly induced under nitrogen limiting conditions (Figure 3A-B).

**Figure 3.**
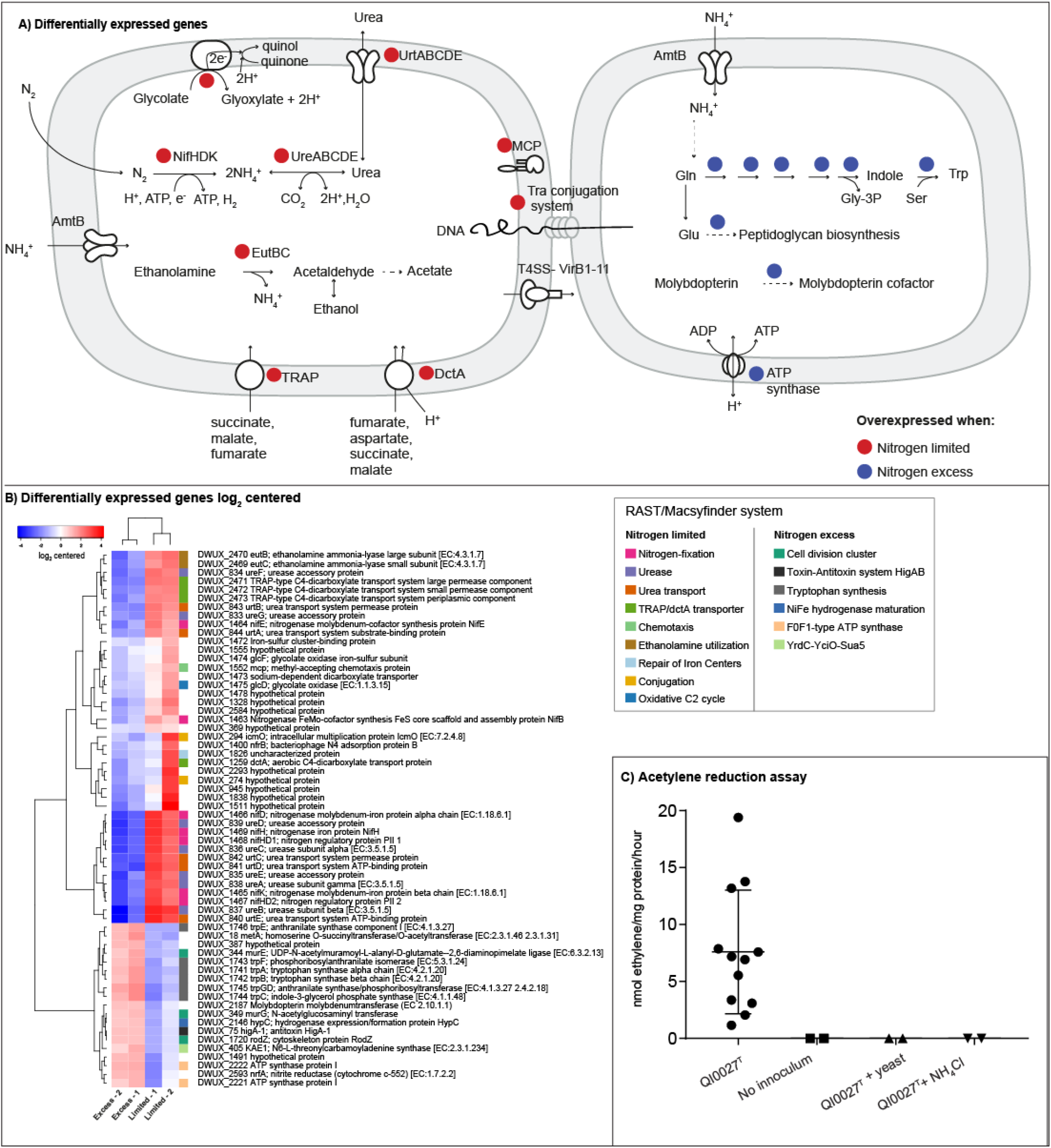
Transcriptome expression and acetylene reduction assay under nitrogen limiting or excess conditions (a) Overview of differentially expressed pathways under nitrogen-limited or excess conditions by QI0027^T^ cultures. Under nitrogen limiting conditions, genes related to nitrogen fixation, urea production and urea transport are overexpressed. Under nitrogen excess conditions, tryptophan synthesis, peptidoglycan biosynthesis and molybdopterin cofactor are overexpressed. p-value cut-off for false discovery rate (FDR) = 0.05; minimum fold change = 3-fold. MCP = methyl-accepting chemotaxis protein; Gln = glutamine; Trp = Tryptophan; Gly-3P = Glyceraldehyde 3 phosphate. Reactions are not balanced. (b) Differentially expressed genes by QI0027^T^ cultures with limiting or excess nitrogen. p-value cut-off for FDR = 0.05; minimum fold change = 3. (c) The strain QI0027^T^ possess nitrogenase activity. Acetylene reduction assay was performed under anaerobic conditions in Hungate tubes as described in materials and methods.

We demonstrated that strain QI0027^T^ was able to fix nitrogen based on an acetylene reduction assay performed in similar growth conditions used for the RNA-seq transcription analysis (Figure 3C). The specific nitrogenase activity rate for strain QI0027^T^ was estimated as 7.6±5.2 nmol ethylene · mg protein^−1^ · h^−1^. As expected, no acetylene reduction was observed when the growth media was supplemented with yeast extract or with 18.54 mM NH_4_Cl (referred as nitrogen excess). To our knowledge, this is the first *Desulfovibrio* isolate from the human gut for which diazotrophy has been physiologically demonstrated.

Further evaluation of our whole transcriptome analysis revealed that the strain QI0027^T^ overexpressed 43 genes under nitrogen limiting conditions, while an additional 19 genes were overexpressed under nitrogen excess conditions (Figure 3B). Under nitrogen excess conditions, QI0027^T^ had a higher expression of genes related to cell cycle and cell division, RNA processing, glycolysis, respiration, as well as cobalamin, ATP, and tryptophan synthesis (Figure 3A-B). Moreover, QI0027^T^ overexpressed the antitoxin HigA, a system that has been linked to survival response during environmental and chemical stresses, including amino acid starvation, growth, and programmed cell death (36).

Besides the nitrogen fixation genes overexpressed under nitrogen limiting conditions discussed above, we observed overexpression of the urease enzyme complex *ureABCEFGD* and the reversible urea uptake system *urtABCDE*. *ureABCEFGD* is used to reversibly transform urea into ammonia and CO_2_. Since QI0027^T^ cultures with limiting nitrogen did not have urea in the media, ammonia resulting from N_2_ fixation was likely converted to urea and the excess excreted. Urea is less toxic than ammonia, more soluble, and can be used as an intermediate molecule for nitrogen excretion (37). Alternatively, some QI0027^T^ cells might have scavenged urea to use it as nitrogen source. Two transporters, from the TRAP and DctA families, which can be used for the import of organic acids (38), were overexpressed under nitrogen limiting conditions, suggesting that efficient transport of dicarboxylates is required to support or regulate nitrogen fixation (39–41).

Interestingly, under nitrogen limiting conditions, QI0027^T^ overexpressed genes that could be used to sense and interact with other cells, including methyl-accepting chemotaxis protein (MCP), the Tra conjugation system and the type IV secretion system (T4SS) when nitrogen was limiting. The Tra conjugation system and T4SS are used by bacteria to exchange genetic material (42, 43). MCP proteins can be used as chemical sensors. Indeed, the MCP gene encoded a protein with two transmembrane domains, a periplasmic binding domain and a cytosolic domain, suggesting that this protein is used to detect external stimulus, possibly the bioavailability of nitrogen (44). Chemotaxis towards chemicals plays a critical role for host colonization by pathogenic and symbiotic bacteria, such as coral pathogens and *Rhizobium* associated to legumes (45). Nitrogen sensing might be also trigger for QI0027^T^ colonization in the gut.

### Prevalence of diazotrophy among *Desulfovibrionaceae*

Several members of the *Desulfovibrionaceae* family possessed the minimal *nif* gene set (*nifH, nifD, nifK, nifE and nifN-B*) for nitrogen fixation but in some cases, these genes may have been lost in several lineages (Figure 1). We searched for the genes related to nitrogen fixation among all available *Desulfovibrionaceae* genomes using tblastn and combined the gene presence/absence with our phylogenomic reconstruction of the *Desulfovibrionaceae* bacteria. Among the lineages that lack the nitrogen-fixing genes, we identified genera that have been linked to disease (e.g. *Bilophila* and *Lawsonia*). Within the ‘*sensu-stricto’ Desulfovibrios*, neither the closed-genome of *D. piger,* nor other almost complete *D. piger* or *D. fairfieldensis* genomes encoded the key gene for nitrogen fixation *nifH*. These *Desulfovibrio* species are the most abundant and wide-spread in western individuals (9). *D. desulfuricans* isolates obtained from free-living environments, as well as our own 13 *D. desulfuricans* isolates obtained from eight Chinese donors, encoded the genes to fix nitrogen. All these *D. desulfuricans* strains grouped closer to *D. legallii* and QI0027^T^.

From all *Desulfovibrionaceae* MAGs recovered from human metagenomes (25), only *D. desulfuricans* and *Desulfovibrio* strains that belong to the same species as QI0027^T^ based on ANI encoded the *nifHBEDK* genes (Figure 1). 12 out of 22 *D. desulfuricans* MAGs, and 16 out of 20 QI0027^T^ MAGs encoded *nif* genes. Since these genomes were recovered from metagenomes, the absence of the gene cluster does not necessarily imply that the gene cluster is not present. To overcome the limitations from binning, we used a targeted assembly approach using the metagenomic reads from the four samples for which the four QI0027^T^ related MAGs did not encode the nitrogenase genes and as reference the QI0027^T^ genome. With this method, we could recover the nitrogenase genes for the four QI0027^T^ MAGs, and based on phylogenetic analysis, the *nifH* gene grouped closer to *nifH* encoded by QI0027^T^. In agreement with our genome searches on the closed genome of *D. piger,* as well as on draft genomes of isolates from *D. fairfieldensis, Bilophila and Lawsonia*, none of the human-derived MAGs from these *Desulfovibrionaceae* species encoded the nitrogenase genes (Figure 1). From the isolates and MAGs that have been sequenced thus far from *‘sensu-stricto’ Desulfovibrios* that encode the genomic repertoire for nitrogen fixation, only *D. legallii,* which was isolated from a shoulder joint infection, *D. desulfuricans* and QI0027^T^ species have been isolated from mammals (Figure 1). The ability to fix nitrogen is therefore not present in the *Desulfovibrionaceae* that are the most abundant in the gut, but rather prevails in *Desulfovibrio* species that tend to be in lower abundance.

### nifH was likely acquired through horizontal gene transfer

The genes required for nitrogen fixation had a patchy distribution among *Desulfovibrionaceae* bacteria. This can be explained by two scenarios that are not mutually exclusive: 1) multiple events of horizontal gene transfer (HGT) have occurred among the *Desulfovibrionaceae* or 2) the gene has been lost multiple times in evolutionary history. To disentangle which of these scenarios was more likely to occur for ‘*sensu-stricto*’ *Desulfovibrio*, we used phylogenetic analysis of the key gene for nitrogen fixation *nifH*. *nifH* has an ancient history of horizontal gene transfer (46). Consistent with previous phylogenetic analyses based on the *nifH* gene product, *nifH* formed four major clades, with clades 1-3 being functional nitrogenases and clade 4 grouping paralogs that are non-functional (46). Most *nifH* sequences from *Desulfovibrionaceae* grouped with *nifH* from cluster 3 (Figure 3). This cluster is known to be present mainly in obligate anaerobes (46). *Desulfovibrionaceae nifH* were not monophyletic and instead formed three major subclades (D1, D2 and D3) (Figure 1). Some *Desulfovibrionaceae* encoded a paralog nitrogenase *anfH* that grouped within cluster 2 (Supplementary Figure 4). Cluster 2 groups alternative nitrogenases that use an Fe-Fe cofactor instead of an Fe-Mo cofactor (46). *nifH* belonging to non *‘sensu-stricto’ Desulfovibrio* grouped within subclades D1 and D2, while ‘*sensu-stricto*’ *Desulfovibrio* bacteria grouped within subclade D3, suggesting at least three independent acquisitions of this gene with multiple gene losses (Figure 1). Subclade D3 is also nested within a group comprising *nifH* from *Clostridia* and *Methanomicrobia* (Archaea), further supporting an independent acquisition of *nifH* by “sensu stricto” *Desulfovibrio* by HGT from *Clostridia*, which was likely to have been more recent compared to D1 and D2 (Figure 1).

## Discussion

We have presented evidence that QI0027^T^ is a novel species with nitrogen-fixing ability distributed in humans from America, Europe and Asia, but with higher representation in Chinese individuals. This novel species was misclassified by commonly used taxonomic classifiers that rely on clade-specific marker genes from metagenomes and could only be identified using whole-genome sequencing and assembly (of MAGs or pure cultures). Although taxonomic classifiers based on clade-specific marker genes claim to be able to distinguish bacteria at species level, these methods rely on proper genus/species classification and characterization, which is problematic among the *Desulfovibrionaceae* as shown by our phylogenomic analyses. Our study combines cultivation, genome sequencing, physiological characterization, differential expression analysis and metagenome sequencing to overcome the bias inherent to each technique.

Based on our acetylene reduction assay and differential expression analyses, strain QI0027^T^ fixed nitrogen only when we did not provide a source of fixed nitrogen in the media, such as yeast extract or NH_4_Cl, which are known to inhibit the metabolically expensive process of nitrogen fixation (47). Our results show therefore that the genes for nitrogen fixation are fully functional in strain QI0027^T^. However, the rate of nitrogen fixation by QI0027^T^ was lower compared to the environmental isolates *D. desulfuricans, D. vulgaris, D. gigas, D. salexigens, D. africanus, and D. thermophilus* (7.6±5.2 vs. 42-918 nmol ethylene · mg protein^−1^ · h^−1^) (11). Postgate and Kent (11) showed with the acetylene reduction assay that 3 out of 15 environmental strains belonging to five *Desulfovibrio* species could not fix nitrogen (11) and hypothesized that the right conditions for nitrogen fixation might have yet to be found. However, this pattern can be explained now as a true lack of the ability to fix nitrogen, since the key gene for nitrogen fixation, *nifH*, has been gained and lost several times in the *Desulfovibrio* clade (Figure 1).

The *nifH* gene present in ‘*sensu stricto*’ *Desulfovibrio* was likely acquired through HGT from a *Clostridia* ancestor. The gut is an environment where *Clostridia* and *Desulfovibrio* frequently encounter each other at high numbers. In this environment, microorganisms are known to rapidly exchange genetic material (48). We hypothesize that *nifH* exchange could have occurred between *Desulfovibrio* and *Clostridia* in an environment where they often encounter each other, such as the gut.

Only a few members of the ‘*sensu-stricto*’ *Desulfovibrio* encoded *nifH. D. desulfuricans* a member of the ‘*sensu-stricto*’ *Desulfovibrio,* is often recovered from human stool samples (3). Some of these *D. desulfuricans* strains might also fix nitrogen, such is the case of our 13 *D. desulfuricans* Chinese gut isolates that encoded the nitrogenase operon. However, not all strains of *D. desulfuricans* encode the genes to fix nitrogen (Figure 1). Indeed, *D. desulfuricans* formed two major clusters with an 80-85% ANI between the two clusters, which shows that these species will need to be reclassified in future studies. The contribution to nitrogen fixation in the human gut by *Desulfovibrionaceae* is therefore likely not limited to QI0027^T^.

### Nitrogen fixation in a nutrient-rich environment

The human gut is a nutrient-rich environment in which most bioavailable nitrogen comes from amino acids present in food. If nitrogen fixation is a high-energy demanding process (49), and the gut is nutrient-rich, which conditions could favour the persistence of *Desulfovibrio* species that can fix nitrogen?

It is commonly assumed that N_2_ fixation can occur only when bioavailable nitrogen is limiting (<1 μM) (50). However, nitrogen fixation can occur in environments with relatively high input of fixed nitrogen such as oceanic waters with high nitrate or ammonia concentrations (30 μM NO_3_-; 200 μM NH_4_ ^+^), symbiotic associations between a diazotrophic cyanobacteria and a single-cell eukaryote living in waters with an excess of nitrogen (UCYN-A/haptophyte symbiosis), as well as the human gut (12-30 mM ammonia in stool, which can be increased with higher protein consumption) (14, 50–53). A human population from Papua New Guinea that had low protein intake (e.g. fixed nitrogen) was shown to have a gut microbiome that was actively fixing nitrogen at higher rates as compared to populations with higher protein intake (49-74 vs. 105 mg/kg body weight/day) based on acetylene reduction assays using stool samples (14, 54). In this human population, nitrogen fixation by the microbiome corresponded to at least 0.01% of the standard nitrogen requirement for humans. Nitrogen fixation was attributed to *Klebsiella* and Clostridia bacteria based on *nifH* sequences recovered from cloning and metagenomic analysis (14). However, the microbial contribution to the nitrogen intake from humans could be population specific.

Some human diets are poorer in protein and hence in bioavailable nitrogen. Until now, nitrogen fixation has not been investigated in Chinese individuals, who are of special interest because their diet tends to be significantly lower in protein content as compared to Mediterraneans, Americans and Japanese (55), although protein content has increased in recent years (56). Nutrition can shape the human microbiome in response to dietary habits. For example, *Bacteroides* from Japanese individuals have acquired porphyranases and agarases through HGT, likely because of the frequent consumption of algae (57). Ammonia is among the metabolites that will have different concentrations depending on diet consumption. Indeed, mice on a diet low in fat and high in plant polysaccharides (LF/HPP) were shown to have lower concentrations of ammonia in the stool compared to mice on a diet high in fat and simple sugars (HF/HS) (~250-300 μM vs. ~350-425 μM) (58). These ammonia concentration differences contributed potentially enough to fitness differences in the high-affinity ammonia assimilation genes for the non-diazotroph *D. piger*. Diazotrophic *Desulfovibrio* species could contribute to the host and the microbiome nitrogen balance, especially when low protein or LF/HPP diets are consumed such is the case of the Chinese diet.

Although the gut may have on average an excess of nitrogen because of the host diet, there may be microniches where bioavailable nitrogen is locally limiting. Therefore, having nitrogen fixation ability would still be a selective advantage. SRB must compete for resources and space to thrive in the gut. Hydrogen, the most commonly used energy source by SRB, is also used by acetogens and methanogens. Niche partitioning could be a mechanism to cope with competitors, as has been observed for even members of the same species (59–61). More than 90% of nitrogen is absorbed in the small intestine and studies using germ-free animals have shown that most gut microbes increase the protein requirement of the host (13, 15). In the large intestine, the bacterial load increases which could create microniches where fixed-nitrogen becomes limiting (62). Reese et al. (15) suggested that the host can secrete nitrogen in the large intestine to control the microbial load and select for preferred microbial taxa based on the comparison of carbon/nitrogen ratios of gut microbes to faeces from 30 mammal species. Under nitrogen limiting conditions, *Desulfovibrio* that can shift their metabolism to fix nitrogen could have an advantage over methanogens, acetogens and non-diazotrophic SRB. Fixed nitrogen could then be used for the synthesis of amino acids, as predicted based on our genomic analyses. Amino acids are a common exchange currency for animals that have established beneficial associations with bacteria that can fix nitrogen, such as termites, clams, sponges and corals (63–65). In rabbits and chicken, bacterial synthesis of amino acids has been shown to be of value to the host (66). Diazotrophic *Desulfovibrio* could use amino acids and ammonia as exchange currency to cooperate with fermenters that release hydrogen. When low protein diets are consumed, the growth of diazotrophic strains could be exacerbated, promoting the cooperation with other members of the microbial milieu, as well as contributing to the nitrogen budget of the host.

An alternative explanation for the presence of diazotrophs in environments with nitrogen excess is that nitrogen fixation is used to maintain an ideal intracellular redox state, analogous to bacteria which can grow heterotrophically but still retain energetically expensive CO_2_ fixation pathways like the Calvin-Benson cycle (67, 68). This has been interpreted as a means to regenerate electron carriers like NAD^+^ under anaerobic conditions, where there can be an excess of reducing equivalents (69). Nitrogen fixation is also energetically expensive and consumes reducing equivalents. So far, the use of N fixation as an electron sink has only been proposed for phototrophs, but perhaps nitrogen fixation could fulfil a similar function in anaerobic niches of the gut, with the additional benefit of producing bioavailable nitrogen. Further work will be required to disentangle which factors trigger nitrogen fixation by *Desulfovibrio* in the human gut.

### Description of *Desulfovibrio diazotrophica* sp. nov

Based on the phylogenomic and physiological data presented in this study, we propose that strain QI0027^T^ is a novel species which we named *Desulfovibrio diazotrophica* sp. nov. QI0027^T^ cells are non-spore forming, Gram-negative rods to spirochete shaped bacteria from the *Deltaproteobacteria* class. Multiplies by binary fission. Cells are 1.4 to 5 μm long and show only sometimes lophotrichous flagella. After 2-7 days at 37°C on Postgate C media, colonies are 0.4-1.9 mm in diameter, pale brown with slightly grey hints to black in colour, round, and with a raised elevation. QI0027 ^T^ grows at 15 to 46°C, (optimum at 37°C), at pH 4.5 to 6.5 (optimum pH 5.9), and with 0-0.51 M NaCl (optimum 0.068-0.171 M NaCl). An obligate anaerobic sulphate reducer that can fix nitrogen. Catalase negative. QI0027 can utilize L-rhamnose, D-fructose, L-fucose, D-galactose, D-galacturonic acid, palatinose, D,L-lactic acid, L-lactic acid, D-lactic acid methyl ester, L-malic acid, pyruvic acid, methyl pyruvate, L-glutamic acid, glucose and D-mannose. The predominant cellular fatty acids are iso C_17:1_ω9c (29%), iso fatty acid methyl ester (C_17:0_ (24.9%) and C_15:0_ (19.3%)), and iso C_17:0_ (17.2-17.5%).

The type strain, QI0027^T^ (=DSMZ 109475 ^T^ = NCIMB 15228^T^ = NCTC 14354^T^), was isolated from a faecal sample donated by a 25-year old male healthy human in Wuxi, China. The G+C content of the type strain is 62.1%.

## Materials and methods

### Isolation

Strain QI0027 ^T^ was isolated from a human stool sample in Wuxi, China in June 2016. The 25-year old male donor was apparently healthy. The donor was provided with a stool collection kit, which included an anaerobic bag, an anaerobic pack (Mitsubishi Gas Chemical, Japan) and ice. The sample was placed inside an anaerobic cabinet within 30 min of collection. A 1 g stool sample was homogenized and used for dilution (ranging from 1:10 to 1:1×10^7^) using anaerobic 1x PBS. Dilutions were plated onto anaerobic Postgate C agar plates, which contained per L of distilled water: 6g sodium lactate; 4.5 g Na_2_SO_4_, 1 g NH_4_Cl, 1 g yeast extract, 0.5 g KH_2_PO_4_, 0.3 g sodium citrate, 0.06 g MgSO_4_.7H_2_O, 0.004 g FeSO_4_.7H_2_O, 4 ml resazurin (0.02% W/V), 0.04 g CaCl_2_.2H_2_O, 0.5 g L-Cysteine hydrochloride, and 15 g agar. pH was adjusted to 7.5±0.1 using 5 M HCl and the media was sterilized by autoclaving. Plates were incubated at 37°C for two weeks. Single black colonies were selected and subcultured two more times until a pure culture of QI0027^T^ was obtained. The purity of the isolate was confirmed by colony morphology, light microscopy, and by checking that all cells in the culture hybridized to the deltaproteobacteria-specific fluorescence *in situ* hybridization probe Delta495a (70) as described by Amann et al., (71). Additionally, 16S rRNA sequencing from colonies that showed slightly different pigmentation or size was done to confirm the purity of QI0027^T^. The 16S rRNA gene was amplified with the AMP_F (5’-GAG AGT TTG ATY CTG GCT CAG) and AMP_R (5’-AAG GAG GTG ATC CAR CCG CA) set of primers according to the method from Baker (72). Purified amplicons were sequenced using the Mix2Seq Kit Overnight service (Eurofins, Germany). To check for the identity of QI0027^T^, the 16S rRNA sequence was compared against the Silva SSU r138 database (73). For long term storage, strain QI0027^T^ was stored at −80°C in Protect Select tubes for anaerobes with cryobeads (TS/73-AN80, Technical Service Consultants, UK).

### DNA extraction, whole genome sequencing and assembly

DNA was extracted using the GenElute Bacterial Genomic DNA kit (NA2100, Sigma-Aldrich, United Kingdom) following the manufacturer’s instructions. For whole-genome sequencing, 947,409 reads were generated using the Illumina HiSeq 3000 platform (Earlham Institute, Norwich, UK). Reads were quality trimmed with bbduk (v.38.06) (sourceforge.net/projects/bbmap) using a minimum quality of 3. Reads were normalized to a maximum coverage of 100 with bbnorm (sourceforge.net/projects/bbmap), and assembled with SPAdes v.3.8.1 (74), which resulted in 100 scaffolds. Quality checks of the genome were done with CheckM (v.1.0.13) (75). The genome was annotated with PATRIC RASTtk (76). Genome statistics were calculated with Quast (v.4.6.1) (73) (shown in Table 1).

### Phylogenomics

Genomes from the *Desulfovibrionaceae* family were obtained from PATRIC (77) and NCBI. Phylogenomic analyses were conducted with the scripts available from phylogenomics-tools (doi:10.5281/zenodo.46122). Conserved marker proteins that are mostly conserved across bacteria were extracted with the Amphora2 pipeline (78). The amino acid sequences encoded by the following 30 marker genes were aligned with Muscle (79): *frr, infC, nusA, pgk, pyrG, rplA, rplB, rplC, rplD, rplE, rplF, rplK, rplL, rplM, rplN, rplP, rplS, rplT, rpmA, rpoB, rpsB, rpsC, rpsE, rpsI, rpsJ, rpsK, rpsM, rpsS, smpB* and *tsf*. Positions of the alignment that were not present in at least 75% of the sequences were removed with Geneious V9 (80). The best evolutionary amino acid substitution model for each marker gene was estimated with RaxML (81). Single amino acid alignments were concatenated to reconstruct a phylogenomic tree with SH-like aLRT support values using RaxML and a partitioned alignment (81). Pairwise average nucleotide identity between the *Desulfovibrionaceae* genomes was estimated with fastANI (82).

### Metagenome sequencing, microbial profiling of stool samples, and detecting presence of QI0027^T^ in Chinese metagenomes

45 humans from China were provided with a stool collection kit as described above. Samples were stored at −80°C upon 48 hours of collection. DNA was extracted from the stool samples using the FastDNA SPIN kit for feces (MP Biomedicals, Shanghai, China) according to the manufacturer’s instructions using approximately 50 mg of stool material per individual. DNA libraries were prepared using the NEB Next Ultra DNA Library Prep kit for Illumina (New England BioLabs). Approximately 10 Gb were sequenced with a HiSeq X Ten instrument as paired-end 150 bp reads. Library preparation and sequencing was done by Novogene (China). Metagenomic reads were quality trimmed with a minimum quality of 2 and human host reads were removed using as reference the human genome GCA_000001405 with bbduk. These clean reads were used to search for QI0027^T^ using three methods: (i) Taxonomic profiling using mOTUs2 (v.2.5.1) with the default database, as well as a modified database that included the genome of QI0027^T^ (23). (ii) Taxonomic profiling using MetaPhlAn2 (v.2.7.7) with the default database because all publicly available human metagenomes have been scanned for the microbial diversity using this tool with default settings (1, 24). (iii) Finally, we mapped the reads against QI0027^T^ genome assembly (only scaffolds >1000 bp) with bbmap (v.38.43) using >95% mapping identity. We considered that QI0027^T^ species was present when > 50% of the reference genome was covered by the reads.

### Recovery of QI0027^T^ from Chinese metagenomes and MAGs

Clean reads of the 45 stool metagenomes were combined and mapped against all publicly available and our own *Desulfovibrionaceae* genomes. The resulting assembly from the combined reads was separated into genomes using an unsupervised binning method based on nucleotide composition, differential coverage and linkage data from paired-end reads (83). Resulting reads were then assembled with metaSPAdes (v.3.13.0) (84) and binned with CONCOCT (v.1.1.0) (83). Genome completeness was estimated using CheckM (v.1.0.13) with the marker lineage *Deltaproteobacteria* (UID3218) (85) and identity was determined with GTDB-Tk (v.1.0.2) using the database release 89 (86). Binning allowed us to recover a total of four *Desulfovibrionaceae* genomes with a completeness ≥97.34% and a contamination ≤8.67%, including QI0027, *Bilophila wadsworthia*, *D. fairfieldensis* and *D. piger*. The relative abundance of reads mapping to the same species as QI0027^T^ was determined using bbmap (v.38.43) as the number of reads mapping to the genome/total number of reads.

The comprehensive compilation of MAGs from human microbiome sequencing efforts by Passolli et al. (1) and Almeida et al. (25) was also exploited to recover *Desulfovibrionaceae* bacteria genomes (opendata.lifebit.ai/table/?project=SGB and ftp.ebi.ac.uk/pub/databases/metagenomics/mgnify_genomes, accessed on March 2020). Metadata for these genomes was retrieved using the *curatedMetagenomicData* R/Bioconductor package (26). To find genomes from the same species as QI0027T, we used ANI similarity of all the Desulfovibrionaceae MAGs against the QI0027^T^ genome with fastANI (82).

### Detection of genes related to nitrogen-fixation and phylogenetic analysis

We searched for the nitrogen fixation related genes among the *Desulfovibrionaceae* genomes included in our phylogenomic tree, as well as the Desulfovibrionaceae MAGs from the collection by Pasolli *et al*., (1). We used as query the amino acid sequences encoded by *nifH, nifD, nifK, nifE* and *nifB* from QI0027^T^ against the isolate genomes and MAGs with tblastn (v.2.2.31+) to be able to detect partial genes at the end of scaffolds (>40% similarity, >40% query coverage, minimum alignment length >50). To improve the genome assembly of the four MAGs that did not encode the nitrogenase cluster, we used read mapping against the QI0027^T^ reference genome using bbmap (v.38.43) with a similarity ≥95% against the reads from projects YSZC12003_3554, SRR3736997, H1M313811and YSZC12003_36012 (Short Read Archive). Results were integrated with the phylogenomic tree with iTol (87). For phylogenetic analysis, we annotated all genomes with Prokka (v.1.14.0) and retrieved *nifH* amino acid sequences. Representative *nifH* amino acid sequences from Clusters 1-4 were retrieved from Gaby and Buckley (46). Sequences were aligned using Mafft v. 1.3.7 and realigned with ClustalW v2.1. The alignment was masked to remove alignment positions with >75% gaps. A maximum-likelihood tree reconstruction was obtained using FastTree (v. 2.1.5) with SH-like branching support values (88).

### Physiology of QI0027^T^

#### pH, salinity, temperature and motility

Physiological testing was done by adjusting Postgate C media without agarose in Hungate tubes (SciQuip, UK) and diluting cells 100-fold. To identify the pH range at which QI0027^T^ could grow, pH was adjusted by injecting deoxygenated 1.34 M HCl for acidic pHs, or 0.16 M NaOH with 10% NaHCO_3_ (w/v) for alkaline pHs before autoclaving. Final pH was measured before inoculation (range from 3.98 to 8.83) and after the growth of the cells. QI0027^T^ grew in media with a pH range between 4.5 to 6.5, and the pH of the media became more basic after cell growth ranging from 6.8 to 7.2. To determine the tolerance of QI0027 ^T^ to different salinity concentrations, growth at 0-0.513 M NaCl was investigated in liquid Postgate C medium as described previously (89). pH and salinity tolerance were investigated at 37°C for 5 days. To test for the effect of temperature on growth, duplicate cultures of strain QI0012 were incubated in 10 ml Hungate tubes at temperature range 10-59°C for up to 7 days. Growth was monitored with a turbidimeter CO 8000 (Biochrom, UK). Motility of QI0027^T^ was tested using semisolid Postgate C agar (5g/L of agar), using as reference for diffusion the motile *D. legallii*. The semisolid media was stabbed with disposable inoculating loops and the diffusion was observed after 3 to 5 days.

#### Antibiotic resistance

Antibiotic resistance was examined by disc diffusion assays on Postgate C agar according to the standard procedure outlined in the National Committee for Clinical Laboratory Standards guidelines (90). Inside an anaerobic cabinet, a sterile swab was dipped in a bacterial cell suspension grown overnight and spread on reduced Postgate C agar media. Antibiotic discs (Oxoid, Thermo Fisher Scientific, UK) were placed on the surface using sterile forceps. Plates were incubated for 48 to 72 h at 37°C. The following antibiotics were tested: ampicillin (25 μg), chloramphenicol (10 μg), nalidixic acid (30 μg), kanamycin (30 μg), erythromycin (30 μg), tetracycline (30 μg), and streptomycin (25 μg).

#### Cell morphology, FISH and transmission electron microscopy

Morphological properties, such as cell shape, cell size and motility were observed by phase-contrast light microscopy (Olympus BX60 microscope). For fluorescence microscopy, a Zeiss Axio Imager M2 microscope was used with a 100x objective lens. Gram staining was done with the Gram staining kit (Sigma-Aldrich, UK) according to the manufacturer instructions. To investigate the presence of flagella, negative staining combined with transmission electron microscopy (TEM) was used. Briefly, a small drop of bacterial cell suspension was applied to a formvar/carbon-coated copper TEM grid (Agar Scientific, Stansted, UK) and left for one minute. Excess liquid was removed with filter paper. A drop of 2% aqueous phosphotungstic acid (PTA) was applied to the grid surface and left for 1 minute. Excess stain was removed with filter paper and the grids were left to dry thoroughly before viewing in the transmission electron microscope. Grids were examined and imaged in a FEI Talos F200C TEM at 200kV with a Gatan One-View digital camera. For scanning electron microscopy (SEM), cells were prepared by critical-point drying (CPD) onto filter membranes (91). Briefly, 100 μL of bacterial cell suspension was added dropwise onto an Isopore membrane polycarbonate filter (HTTP01300, Millipore, UK). The filter was trimmed beforehand with a razor blade to identify the surface with inoculum. Cells were left to adhere to the surface for 5 minutes after which the filters were placed in 2.5% glutaraldehyde in 0.1 M PIPES buffer (pH 7.2) and fixed for 1 hour. After washing with 0.1 M PIPES buffer, each sample was carefully inserted into a CPD capsule and dehydrated in a series of ethanol solutions (30, 50, 70, 80, 90, 3x 100% including a final dry ethanol change). Samples were dried in a Leica EM CPD300 Critical Point Dryer using liquid carbon dioxide as the transition fluid. The filters were carefully mounted onto SEM stubs using sticky tabs, ensuring that the inoculated surface was facing upwards. The samples were coated with platinum in an Agar high-resolution sputter-coater apparatus. Scanning electron microscopy was carried out using a Zeiss Supra 55 VP FEG SEM, operating at 3kV.

#### Fatty acid analyses

For determination of cellular fatty acids from the strains *D. legallii* (DSM 19129) and QI0027^T^, cells were grown in Postgate C media at 37°C for 18 h. Cells in the exponential phase were pelleted by centrifugation at 4,000 g for 10 min. Fatty acid identification was carried out by the Identification Service of the DSMZ (Braunschweig, Germany). Fatty acids were converted to fatty acid methyl esters (FAMEs) by saponification, methylation and extraction according to Miller (92) and Kuykendall et al., (93). Samples were prepared according to the standard method given in MIDI Technical Note 101. Profiles of fatty acids were determined with an Agilent 6890N gas chromatograph and processed with the MIDI Inc Sherlock MIS software version 6.1.

#### Utilization of carbon substrates

To determine which carbon substrates could be utilized by QI0027^T^ and *D. legallii*, we used AN microplates from the OmniLog™ system (1007, Technopath, Ireland). We followed the manufacturer’s instructions (AN Microplate™ procedure for testing carbon metabolism of anaerobic bacteria, draft of 3.11.2017 and (94)). Briefly, cells of the two species were grown on Biolog Universal Anaerobe agar (Technopath, Ireland) inside an anaerobic cabinet until visible growth could be observed (2-3 days). Cells were collected with cotton swabs and used for preparing a cell suspension in AN inoculating fluid (Technopath, Ireland) with turbidity between 0.4 to 0.48. These suspensions were used for the inoculation of the AN microplate in triplicates. To maintain an anaerobic atmosphere during the experiment, AN microplates were placed inside plastic bags which had an atmosphere of 90% N_2_, 5% CO_2_ and 5% H_2_. The plates were incubated at 37°C for 36 h and the colour change caused by the response to the oxidation of the different substrates in the plate were measured with an Omnilog plate reader (Biolog). The resulting data were analysed with the opm package (v.1.3.63) (95) implemented in R. Reactions were defined as positive or negative using k-means partitioning as implemented in the function do_disc.

### Determination of nitrogenase activity

To determine the nitrogenase activity of QI0027^T^ to fix nitrogen, we used the acetylene reduction assay (ARA) (96, 97). The cells were first grown in media free of added fixed nitrogen (ammonia, NH_3_), modified from Postgate B media (98), which contained per L of distilled water: 3.5 g sodium lactate, 2 g MgSO_4_.7H_2_O, 0.25 g CaSO_4_, 0.5 g K_2_HPO_4_, 0.004 g FeSO_4_.7H_2_O, 9 g NaCl, 4 ml resazurin (0.02% W/V), 0.1 g ascorbic acid, and 0.1g thioglycolic acid. pH was adjusted to 7.5±0.1 using 5 M HCl. The media was dispensed into Hungate tubes (6.2 mL gas phase) and sterilized by autoclaving. Media was supplemented with 1 ml/L of trace element solution SL-10 (DSMZ). Growth could be observed after 5 to 7 days. These cells were used to inoculate 12 tubes with a 1:100 dilution in 10 ml of media. As nitrogenase activity is inhibited when fixed nitrogen is available, we included as negative control duplicate tubes supplemented with either 1g/L of yeast extract or 1g/L NH_4_Cl (47, 98, 99). Duplicate blank tubes without inoculum were also included as negative controls. All tubes were injected 10% v/v of the gas phase with acetylene and incubated at 37°C. After 7 days incubation, ethylene formation was quantified by using a Perkin Elmer Clarus 480 gas chromatograph equipped with a HayeSep® N (80-100 MESH) column. The injector and oven temperatures were kept at 100 °C, while the FID detector was set at 150 °C. The carrier gas flow was set at 8 - 10 mL/min. The ethylene standard was prepared from ethephon decomposition based on Zhang and Wen (100). Nitrogenase activity is reported as nmol of ethylene produced per mg protein per hour. To determine whole-cell protein concentration, 5 ml of the culture was centrifuged at 3,200 g for 10 min at 4°C. The pellet was resuspended in 500 μl NaOH 0.1 M and incubated for 3 h at 30°C. Cells were then sonicated using an MSE Soniprep 150 instrument for seven cycles of 15 s sonication followed by at least 30 s rest on ice. Protein content from the lysates was finally measured with the Bradford assay (Biorad).

### Transcriptional response to presence and absence of ammonia

To identify the genes that are expressed by QI0027^T^ when fixing nitrogen, we used whole transcriptome sequencing of cultures grown in modified Postgate B minimal media without any source of fixed nitrogen or with 1 g/L NH_4_CL. QI0027^T^ cells grown for 48 h in Postgate C media were centrifuged at 4,000 g for 3 min. Cells were resuspended in modified Postgate B media without NH_4_CL. These cells were used to inoculate six Hungate tubes with 10 ml Postgate B media, six tubes supplemented with 1 g/L NH_4_Cl, as well as duplicate control tubes supplemented with 1g/L of yeast extract using a 1:100 dilution. Duplicate negative controls of Postgate B media supplemented with yeast extract without QI0027^T^ inoculum were also included. As expected, control tubes with yeast extract showed growth in 48 h. After 7 days, we divided the media from six tubes per condition into two to increase the input biomass for RNA extraction. Cells were centrifuged at 4,000 g for 10 min and immediately frozen in dry ice for 10 min. RNA was extracted using the RNeasy Plus Minikit (Qiagen, UK) according to the manufacturer’s instructions with the following modifications. 350 μl RNeasy lysis buffer (RLT) and 10 μl ß-mercaptoethanol were added to the cell pellets. Cells were disrupted in a FastPrep (MP Bio, UK) with 3 cycles lasting 20 sec at 4 m/s in Lysing Matrix E tubes (MP Bio, UK). Samples were centrifuged at 13,000 g for 3 min, and the supernatant was used for further RNA extraction. Resulting RNA was used for library construction using the TruSeq Total RNA (NEB Microbe) kit without rRNA depletion. 3 Gb per sample were generated as 100bp paired-end reads using a NovaSeq instrument at Macrogen (South Korea).

For differential expression analyses, adapters and rRNA sequences were removed from transcriptome reads using bbduk (v.38.06). Clean reads were mapped to the QI0027^T^ reference genome with a minimum identity of 95% and all ambiguous mapping reported. The number of transcripts per gene was estimated with featureCounts (v.2.0) (101). Differentially expressed genes were detected with edgeR with trimmed mean of M values (TMM) normalization (102, 103). Pathway differentially expressed were examined with Pathway Tools (v.23.5). Operons were predicted with Rockhopper. Secretion and conjugation systems were predicted with Maccsyfinder (v.1.0.5) using the CONJScan and TXSS models (104, 105). Transmembrane domains of the MCP protein was predicted with TMHMM (v.1.0.1).

## Data availability

All sequencing data was submitted to the European Nucleotide Archive (http://www.ebi.ac.uk/ena/data/view/). Sequencing reads and the genome assembly of QI0027^T^ are available with the accession number PRJEB34368. Metagenomic reads obtained from Chinese stool samples without human reads are available with the accession number PRJEB38832. Transcriptomic reads are available under the accession number PRJEB38833.

## Author contributions

LS and AN designed the study. LS conducted experiments and analysed data. TL isolated SRB, collected stool samples and extracted nucleic acids for metagenomes. MBB and LS did acetylene reduction assays. BKBS suggested the impact of nitrogen fixation as electron sink and critically commented on the manuscript. CB did electron microscopy. QZ and CW coordinated sample collection in China. LS wrote the manuscript with contributions from all co-authors.

## Ethics statement

This study was approved by the Ethics Committee in Jiangnan University, China (SYXK 2012-0002). All the faecal samples from healthy persons were for public health purposes and these were the only human materials used in present study. Written informed consent for the use of their faecal samples was obtained from the participants or their legal guardians. All of them conducted health questionnaires before sampling and no human experiments were involved. The collection of faecal samples had no risk of predictable harm or discomfort to the participants.

## Acknowledgements

We thank Gemma Langridge, Claire Hill and Barry Bochner for technical support using the Omnilog machine. We thank the JIC Bioimaging facility for access to electron microscopes.

---

**Supplementary Figure 1:**
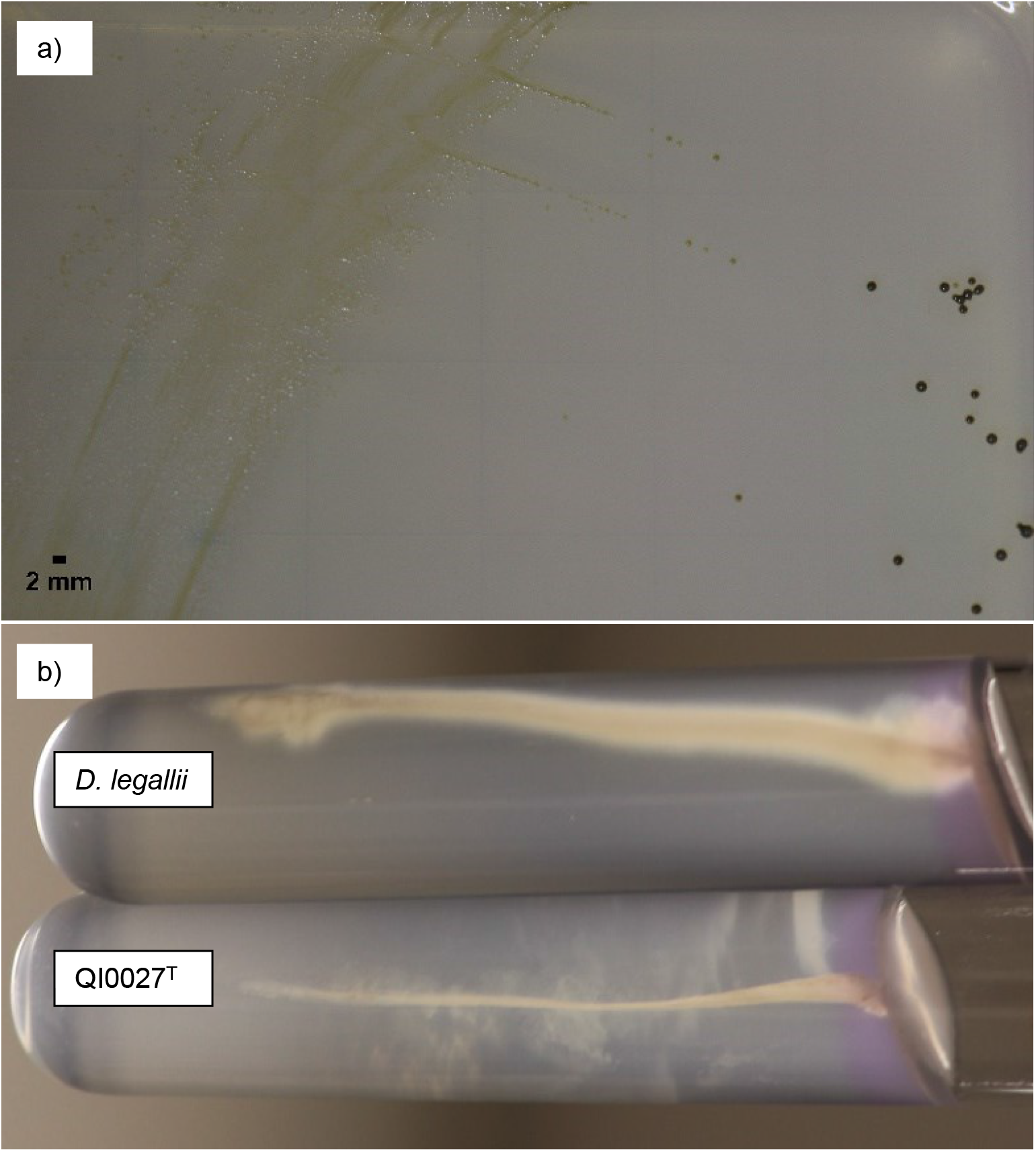
Colony morphology of QI0027^T^ (a) and motility assay (b). For the motility assay, soft agar was stabbed with an inoculum of QI0027 or *D. legallii* anaerobically. Cells were grown in Postgate C media with 5 g/L agarose for 72 hrs.

**Supplementary Figure 2:**
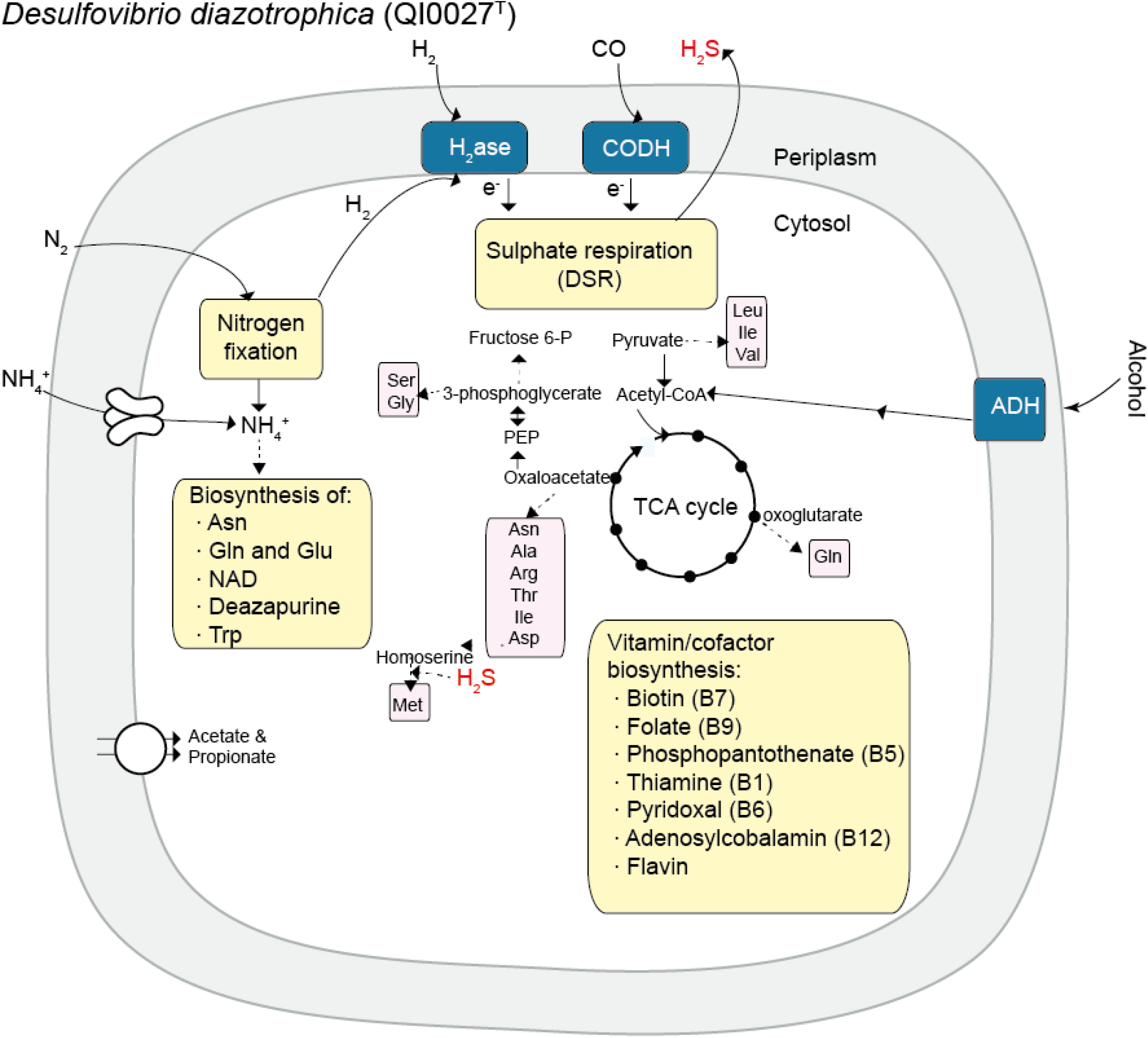
Metabolic reconstruction of QI0027^T^ based on genome information. H_2_ase = hydrogenase; ADH = alcohol dehydrogenase; CODH = carbon monoxide dehydrogenase.

**Supplementary Figure 3:**
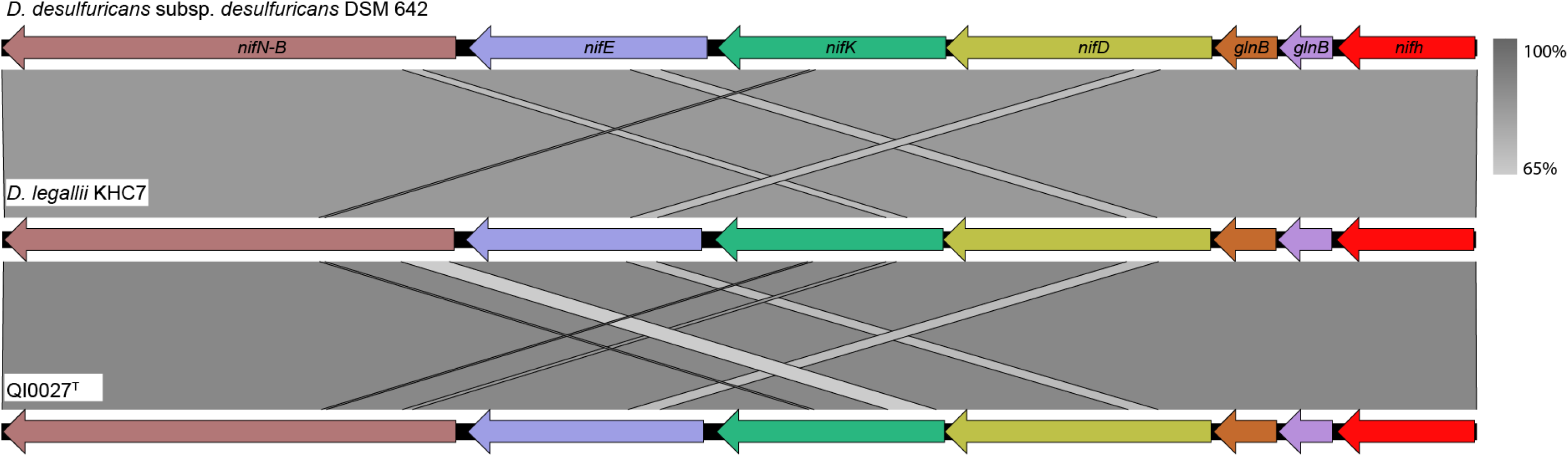
The *nif* gene cluster is highly conserved among QI0027^T^, *D. legallii* and *D. desulfuricans* subsp. desulfuricans DSM 642. Similarity between genomic regions was estimated with blastn incorporated into Easyfig (106).

**Supplementary Figure 4:**
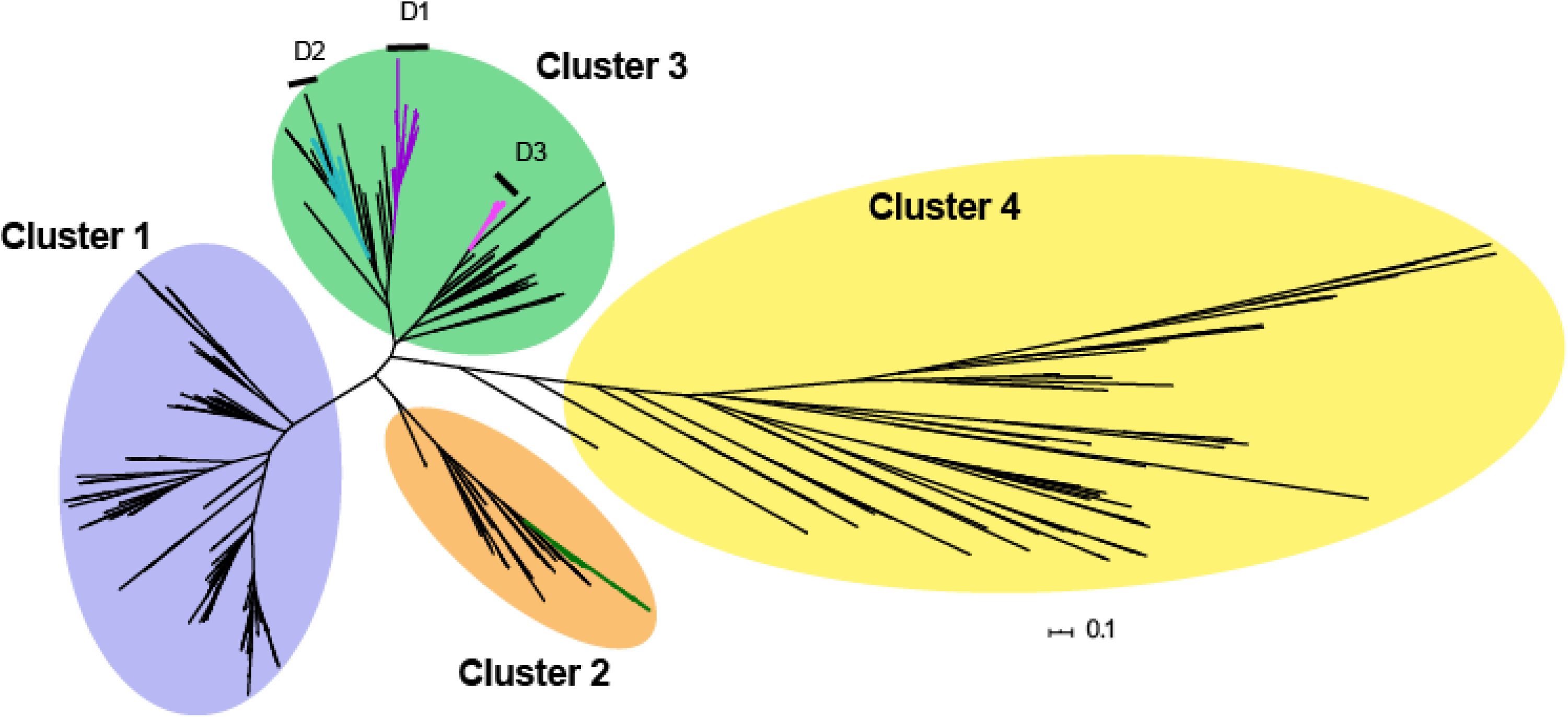
Overview of *nifH* phylogeny shown in Figure 1, unrooted. Some Desulfovibrionaceae have an additional *anfH* nitrogenase that falls within Cluster 2. Cluster 2 genes are paralog genes that can also fix nitrogen using an Fe-Fe cofactor.

**Supplementary data set 1:** Relative abundance of QI0027^T^-related species in 45 Chinese stool metagenomes. The abundance was estimated based on read mapping against QI0027^T^ scaffolds ≥1000 bp. The relative abundance of the two *Desulfovibrionaceae* species predicted by MetaPhlan2 is also shown.

**Supplementary data set 2:** Differential gene expression analysis of QI0027^T^ cultures with limiting or excess nitrogen. Differentially expressed (DE) genes were identified using edgeR. Fold change expression values were estimated using (log_2_(expression with nitrogen excess)-log_2_(expression with nitrogen limitation)). Thus, negative log_2_ fold change values indicate genes overexpressed under nitrogen limiting conditions, while positive values indicate genes overexpressed under excess conditions. Read counts were normalized with TMM. For clustering analysis (Figure 3B), log_2_ centred values were estimated from the TMM normalized values. Genes overexpressed with nitrogen deplete conditions are shown in blue. Genes overexpressed with nitrogen replete conditions are shown in orange. **p-value significant corrected using false discovery rate (FDR) correction. *Based on FDR non-significantly DE, but gene operon had non-corrected p-value <0.05. DE = Differentially expressed.

## Abbreviations

ANI: average nucleotide identity
FAME: fatty acid methyl ester
*D*.: *Desulfovibrio*
SRB: sulphate reducing bacteria
TEM: transmission electron microscopy
SEM: scanning electron microscopy
MAGs: metagenome-assembled genomes
WGS: whole genome sequencing
T4SS: type IV secretion system
MCP: methyl-accepting chemotaxis protein

